# Sweet taste of heavy water

**DOI:** 10.1101/2020.05.22.110205

**Authors:** Natalie Ben Abu, Philip E. Mason, Hadar Klein, Nitzan Dubovski, Yaron Ben Shoshan-Galeczki, Einav Malach, Veronika Pražienková, Lenka Maletínská, Carmelo Tempra, Victor Cruces Chamorro, Josef Cvačka, Maik Behrens, Masha Y. Niv, Pavel Jungwirth

**Author notes:** Both authors contributed equally (Natalie Ben Abu, Philip E. Mason).

## Abstract

Hydrogen to deuterium isotopic substitution has only a minor effect on physical and chemical properties of water and, as such, is not supposed to influence its neutral taste. Here we conclusively demonstrate that humans are, nevertheless, able to distinguish D_2_O from H_2_O by taste. Indeed, highly purified heavy water has a distinctly sweeter taste than same-purity normal water and adds to perceived sweetness of sweeteners. In contrast, mice do not prefer D_2_O over H_2_O, indicating that they are not likely to perceive heavy water as sweet. HEK 293T cells transfected with the TAS1R2/TAS1R3 heterodimer and chimeric G-proteins are activated by D_2_O but not by H_2_O. Lactisole, which is a known sweetness inhibitor acting via the TAS1R3 monomer of the TAS1R2/TAS1R3, suppresses the sweetness of D_2_O in human sensory tests, as well as the calcium release elicited by D_2_O in sweet taste receptor-expressing cells. The present multifaceted experimental study, complemented by homology modelling and molecular dynamics simulations, resolves a long-standing controversy about the taste of heavy water, shows that its sweet taste is mediated by the human TAS1R2/TAS1R3 taste receptor, and opens way to future studies of the detailed mechanism of action.

**One sentence summary:** Heavy water elicits sweet taste for humans via the TAS1R2/TAS1R3 taste receptor.

## Main

Heavy water, D_2_O, has fascinated researchers since the discovery of deuterium by Urey in 1931(*1, 2*). The most notable difference in physical properties between D_2_O and H_2_O is the roughly 10% higher density of the former liquid, which is mostly a trivial consequence of deuterium being about twice as heavy as hydrogen. A more subtle effect of deuteration is the formation of slightly stronger hydrogen (or deuterium) bonds in D_2_O as compared to H_2_O(*3, 4*). This results in a small increase of the freezing and boiling points by 3.8°C and 1.4°C, respectively, and in a slight increase of 0.44 in pH (or pD) of pure water upon deuteration(*5*). In comparison, a mere dissolution of atmospheric CO2 and subsequent formation of dilute carbonic acid in open containers has a significantly stronger influence on the pH of water, changing it by more than one unit(*6*).

Biological effects are observable for high doses of D_2_O. While bacteria or yeasts can function in practically pure D_2_O, albeit with somewhat hindered growth *rate*(*7–9*), for higher organisms damaging effects on cell division and general metabolism occur at around 25% deuteration, with lethal conditions for plants and animals typically occurring at ~40-50% deuteration of the body water (*2, 10, 11*). Small levels of deuteration are, nevertheless, harmless. This is understandable given the fact that about 1 in every 6400 hydrogens in nature is found in its stable isotope form of deuterium(*12*). Oral doses of several milliliters of D_2_O are safe for humans(*13*) and are being routinely used together with another stable isotopic form of water, H2^18^O, for metabolic measurements in clinical praxis(*14*). Probably the best-established effect of D_2_O is the increase of the circadian oscillation length upon its administration to both animals and plants. This has been attributed to a general slowdown of metabolism upon deuteration, although the exact mechanism of this effect is unknown(*15, 16*).

A long-standing unresolved puzzle concerns the taste of heavy water. There is anecdotal evidence from the 1930s that the taste of pure D_2_O is distinct from the neutral one of pure H_2_O, being described mostly as “sweet”(*17*). However, Urey and Failla addressed this question in a short article published in Science in 1935 concluding authoritatively that upon tasting “neither of us could detect the slightest difference between the taste of ordinary distilled water and the taste of pure heavy water”(*18*). This had, with a rare exception(*19*), an inhibitive effect on further human studies, with research concerning effects of D_2_O focusing primarily on animal or cell models. Experiments in animals indicated that rats developed aversion toward D_2_O when deuteration of their body water reached harmful levels, but there is conflicting evidence to whether they can taste heavy water or use other cues to avoid it(*20, 21*).

Within the last two decades, the heterodimer of the taste receptor of the TAS1Rs type of G-protein coupled receptors (GPCRs), denoted as TAS1R2/TAS1R3, was established as the main receptor for sweet taste(*22*). The human TAS1R2/TAS1R3 heterodimer recognizes diverse natural and synthetic sweeteners(*23*). The binding sites of the different types of sweeteners include an orthosteric site (a sugar-binding site in the extracellular Venus flytrap domain of TAS1R2) and several allosteric sites, including sites in the extracellular regions of the TAS1R2 and TAS1R3 subunits and in the transmembrane domain of TAS1R3(*24, 25*) (Figure 1). Additional pathways for sweet taste recognition have also been suggested, involving glucose transporters and ATP-gated K+ channel(*26, 27*).

**Figure 1.**
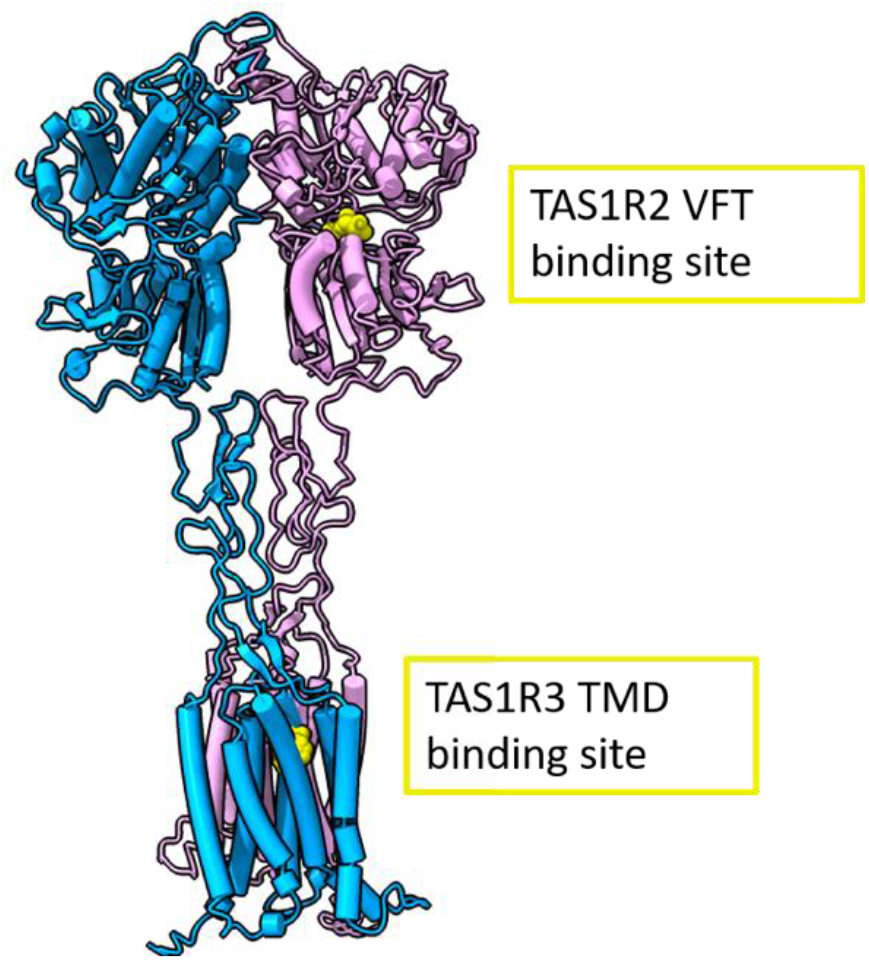
Full human sweet taste TAS1R2/TAS1R3 receptor model with the TAS1R2 monomer colored in pink and the TAS1R3 monomer in cyan. Binding sites are represented in yellow. The full receptor heterodimer was prepared with the I-Tasser web server(*31*) based on multiple published structures (i.e. 6N51, 5X2M, and 5K5S). The structures were aligned to a Metabotropic Glutamate Receptor 5 Cryo-EM structure (PDB: 6N51) and minimized using Schrödinger Maestro 2019-1. The binding site of TAS1R2 is based on coordinates of docked D-glucose to a Venus flytrap (VFT) model that was previously validated (*32*) (modeling based on template PDB ID: 5X2M, docking performed with Schrödinger Maestro 2019-1, Glide XP), and the TAS1R3 binding site is based on a lactisole molecule docked to the TAS1R3 TMD model (template PDB IDs: 4OR2 and 4OO9, Schrödinger Maestro 2018-2, Glide SP). The figure was made using ChimeraX (version 0.93)(*33*).

Interestingly, not all artificial sweeteners are recognized by rodents(*28*). Differences in human and rodent responses to tastants, as well as sweetness inhibitors such as lactisole, have been useful for delineating the molecular recognition mechanism of sweet compounds – using human-mouse chimeric receptors, it was shown that the transmembrane domain (TMD) of human TAS1R3 is required for the activating effects of cyclamate(*29*) and for the inhibitory effect of lactisole(*30*).

A combination of TAS1R3 with another member of the TAS1R family, TAS1R1, results in a dimer that mediates umami taste, elicited by molecules such as glutamate or its sodium salt form of monosodium glutamate (MSG)(*34*). Bitter taste is mediated by the taste receptors type 2 (TAS2R) gene family(*35*), a branch of Family A GPCRs(*24*). The human genome has 25 TAS2R subtypes and over a thousand of bitter compounds are currently known(*36*), with numerous additional bitter tastants predicted (*37*).

In this study, we systematically address the question of sweet taste of heavy water by a combination of sensory experiments in humans, behavioral experiments in mice, tests on sweet taste receptor-transfected cell lines, and computational modeling including molecular dynamics (MD) simulations. This combined approach consistently leads to a conclusion that the sweet taste of pure D_2_O is a real effect for human subjects due to activation of the TAS1R2/TAS1R3 sweet taste receptor. While present simulations show, in accord with previous experiments(*38*), that proteins are systematically slightly more rigid and compact in D_2_O than in H_2_O, the specific molecular mechanism of the heavy water effect on the TAS1R2/TAS1R3 receptor remains to be established.

## Water purity

We have paid great attention to the purity of the water samples, further degassing and redistilling under vacuum the purest commercially available D_2_O and H_2_O. The lack of non-negligible amounts of organic impurities was subsequently confirmed by gas chromatography with mass spectrometry analysis and by experiments with water samples at different levels of purification, see Supporting Material (SM). This is extremely important – note in this context that “the vibrational theory of olfaction”, which suggested distinct perception of deuterium isotopes of odorants due to difference in their vibrational spectra(*39*), has been refuted with some of the observed effects turning out to be due to impurities(*40, 41*).

## Experiments with a human sensory panel

A human sensory panel was employed to study the D_2_O taste. Triangle tests based on two samples of H_2_O and one sample of D_2_O (or vice versa), with random success rate of one third, were presented to the panelists in a randomized order. Panelists were asked to pick the odd sample out - to smell only, to taste only (with a nose clips), or to taste with open nose. Our results show that humans perceive D_2_O as being clearly distinguishable from H_2_O based on its taste: In open nose taste test 22 out of 28 participants identified the odd sample correctly (p=0.001), and in taste only test 14 out of 26 identified the odd sample correctly (p=0.03). However, in smell-only triangle test, only 9 out of 25 panelists chose the odd sample correctly (p>0.05). Data are summarized in Figure S1A-C in SM.

Next, the perceived sweetness of D_2_O in increasing proportion to H_2_O was reported using a 9-point scale. D_2_O sweetness was shown to increase in a dose-dependent manner (Figure 2A). The perceived sweetness of increasing concentrations of glucose (Figure 2B), sucrose (Figure 2C), and an artificial sweetener cyclamate (Figure 2D) was tested when dissolved in H_2_O or in D_2_O. Pure D_2_O was again perceived as slightly sweet and significantly sweeter than H_2_O. Furthermore, D_2_O adds to the sweetness of glucose, cyclamate, and low concentrations of sucrose.

**Figure 2.**
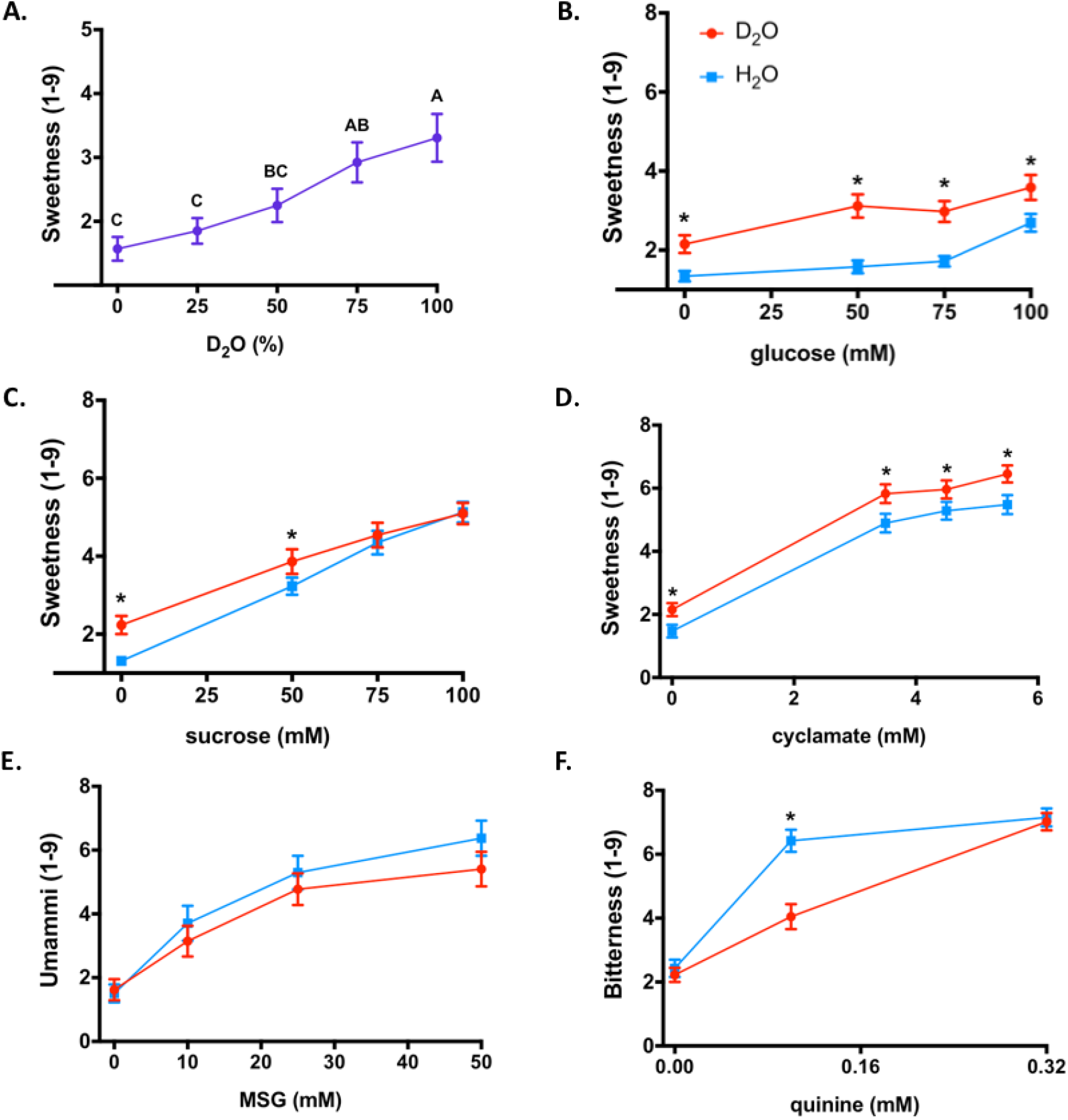
D_2_O sweetness and its effect on different tastants. (A) Sweetness of D_2_O mixed at increasing ratios with H_2_O. Treatments not connected by the same letters are significantly different (p<0.05 in Tuckey Kramer test). (B)-(F) The effect of D_2_O (red) compared to H_2_O (blue) on glucose (B), sucrose (C), cyclamate (D), quinine (E), and MSG (F) taste-specific intensity. Asterisks indicate a significant (p < 0.05) difference between water types using the two-way analysis of variance (ANOVA) with a pre-planned comparison t-test. All data are presented as the mean ± the Standard Error of Measurement (SEM); n=15-30 (4-12 males). The y axis shows the response for individual modalities, while the x axis is labeled with different water samples. Scale for each modality is labeled as 1 = no sensation, 3 = slight, 5 = moderate, 7 = very much, and 9 = extreme sensation.

We then checked whether the effect of D_2_O is sweetness-specific or general, whereby D_2_O might also add intensity to savory taste of umami compounds (MSG) and to bitter taste of bitter compounds (quinine), since these taste modalities are also mediated by GPCR receptors expressed in taste cells. We established here that the intensity of savory taste of MSG in D_2_O did not differ from that in H_2_O (Figure 2E), while the perceived bitterness of quinine was in fact slightly reduced in D_2_O compared to quinine in H_2_O (Figure 2F). This is in agreement with the known effect of sweeteners as maskers of bitter taste, that may be due to both local interactions and sensory integration effects(*42–44*). Thus, we have ascertained that D_2_O is sweet and adds to sweetness of other sweet molecules, but not to intensity of other GPCR-mediated taste modalities.

## Experiments with mice

Next, we addressed the question whether the sweetness of D_2_O is perceived also by rodents. Lean mice of the C57BL/6J strain were drinking pure H_2_O, D_2_O, or a 43 mmol/l H_2_O sucrose solution for 16h during a night period. Namely, each of the three groups of mice had a choice from two bottles containing i) H_2_O and D_2_O, ii) H_2_O and sucrose solution, or iii) H_2_O and H_2_O (as a control). The food intake was unaffected in all groups (see SM).

The results of the drinking experiments are presented in Figures 3A-C, with a snapshot of the experimental setup shown in Figure 3D. In cages where mice were offered both normal water and heavy water (Figure 3A) consumption of D_2_O was within statistical error the same as that of H_2_O. Previous reports have shown that on longer timescales than those reported here mice learned to avoid D_2_O, as it is poisonous to them in larger quantities(*10*). It is not clear what is the cue that enables the avoidance learning, but it is evident that the early response to D_2_O is not attractive, suggesting that it is not eliciting sweet taste in mice.

**Figure 3.**
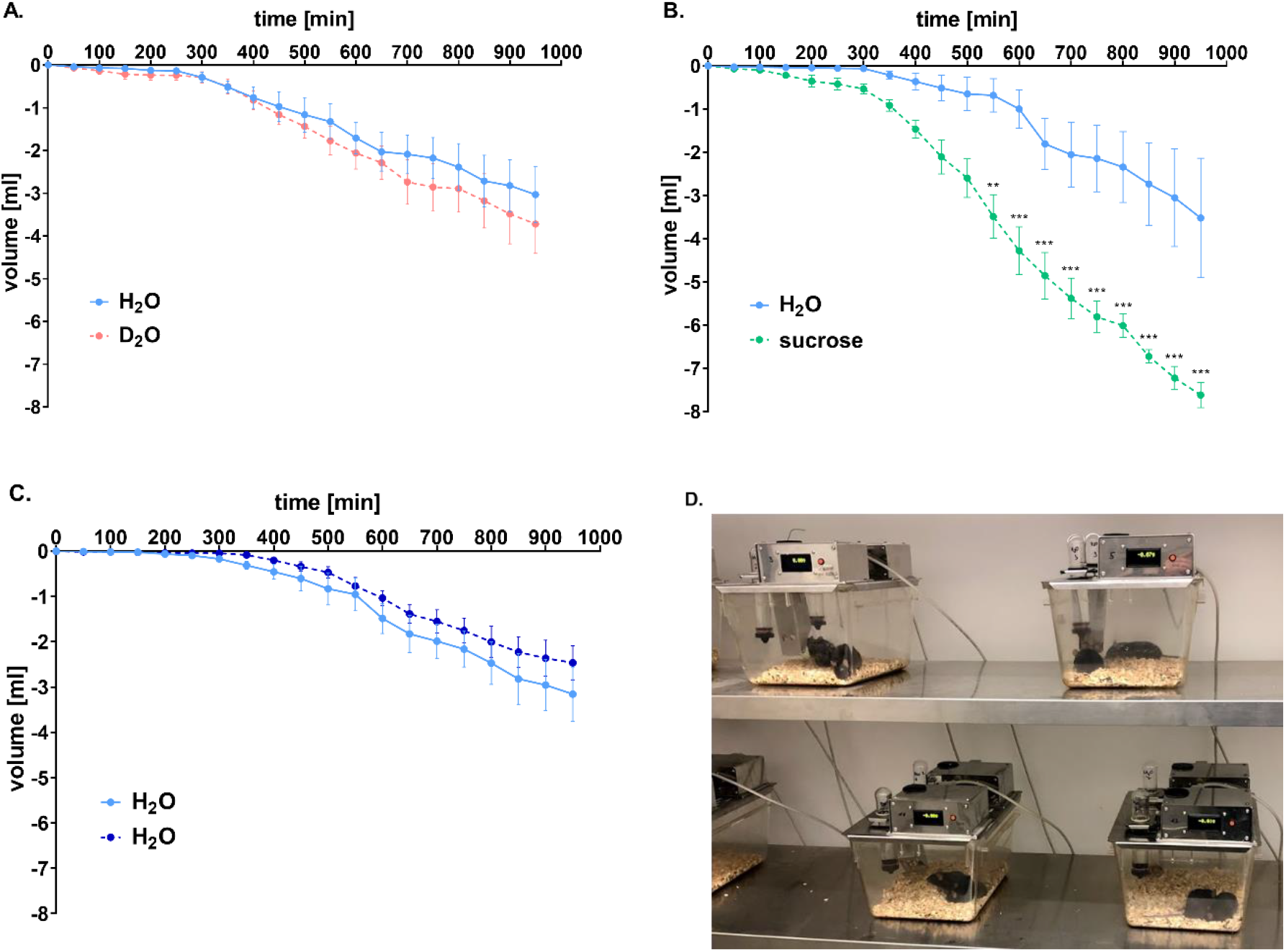
Time-resolved volumes of water consumption by mice. (A) Volume consumption of D_2_O is not different from that of H_2_O (n = 12). (B) Mice show strong preference to sucrose solution (n = 10). Significance is ** p < 0.01, *** p < 0.001. (C) Volume consumption of the control group drinking H_2_O only (n = 12). (D) Snapshot of the automatic drinking monitoring system. Mice were placed in groups of two in individual cages. Data are presented as the mean ± standard error of the mean (SEM). Statistical analysis was performed using two-way ANOVA with a Bonferroni’s multiple comparisons test.

By contrast, mice exhibit a strong preference for sucrose solution over H_2_O. Indeed, the consumed volume was significantly increased in line with the predilection of mice for sucrose solutions (Figure 3B). The amount of H_2_O consumed by the control group from either of the two bottles, both containing H_2_O, is depicted in Figure 3C. Overall, the data shows that in all three experiments mice consumed comparable amounts of H_2_O and D_2_O, with significant increase of consumption of the sucrose solution.

## Assessing involvement of TAS1R2/TAS1R3 receptor using human sensory panel

The chemical dissimilarity of D_2_O from sugars and other sweeteners raises the question whether the effect we observed in human subjects is mediated by TAS1R2/TAS1R3, which is the major receptor for sweet taste(*22*). This was first explored by combining water samples with lactisole as an established TAS1R2/TAS1R3 inhibitor(*30*). Using the two-alternative forced choice (2AFC) method, in which the participant must choose between two samples, 18 out of 25 panelists chose pure D_2_O as sweeter than D_2_O + 0.9 mM lactisole solution (p<0.05, Figure 4A). In an additional experiment, the sweetness of pure D_2_O was scored significantly higher than that of D_2_O + 0.9 mM lactisole solution (p=0.0003), while the same amount of lactisole had no effect on the perception of sweetness of H_2_O that served as control (Figure 4B). These results suggest that D_2_O elicits sweetness via the TAS1R2/TAS1R3 sweet taste receptor.

**Figure 4.**
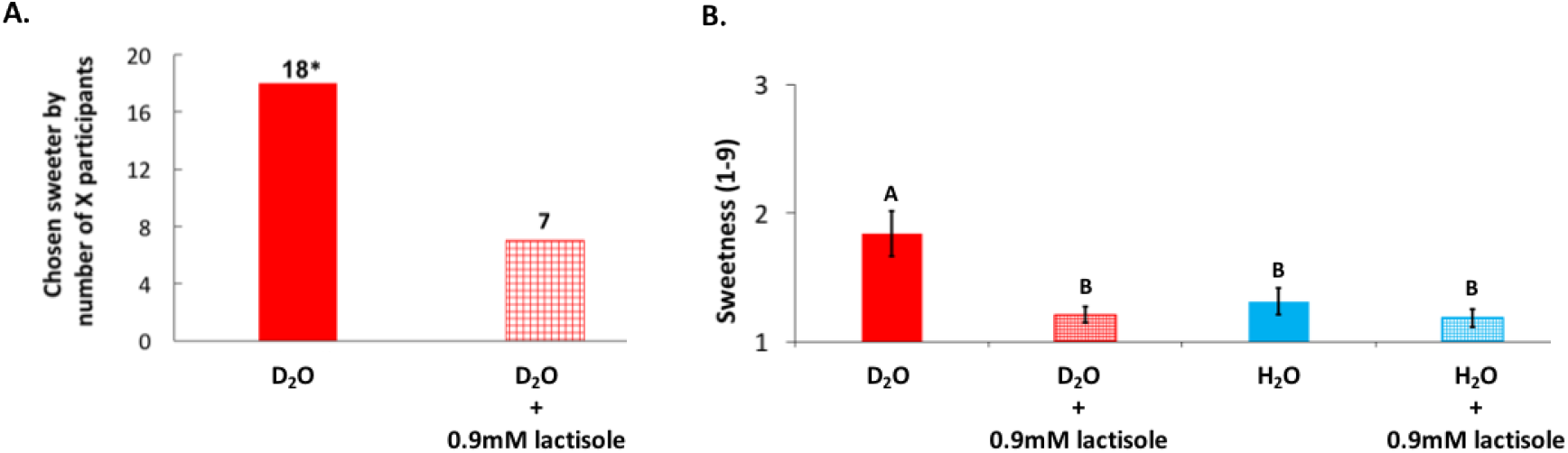
Lactisole reduces sweetness of D_2_O. (A) 2AFC test. Pure D_2_O was chosen to be sweeter (p<0.05) than the sample with lactisole by 18 participants (n=25; 11 males). (B) Effect of 0.9mM lactisole on sweetness intensity using the 9-point scale. Data are presented as the mean ± SEM. The y axis shows the response for sweetness on a 9-point scale, while the x axis is labeled with different water samples. Statistical analysis was performed using ANOVA with a Tuckey Kramer test (n=27; 9 males); treatments not connected by the same letters are significantly different (p<0.05). Scale for sweetness is labeled as 1 = no sensation, 3 = slight, 5 = moderate, 7 = very much, and 9 = extreme sensation.

## Cell-based experiments for establishing the role of TAS1R2/TAS1R3

To confirm the involvement of the sweet taste receptor TAS1R2/TAS1R3 in D_2_O signaling we performed functional calcium mobilization assays using HEK 293 FlpIn T-Rex cells heterologously expressing both required TAS1R subunits as well as the chimeric G protein Gα15Gi3(*45, 46*). As seen in Figure 5, D_2_O at 1.85 M and 5.84 M concentrations in H_2_O (3.3 % and 10.4 % respectively) elicited robust responses in TAS1R2/TAS1R3 expressing cells. The strong reduction or absence of D_2_O-elicited fluorescence response in the presence of lactisole confirmed the dependence on TAS1R2/TAS1R3.

**Figure 5.**
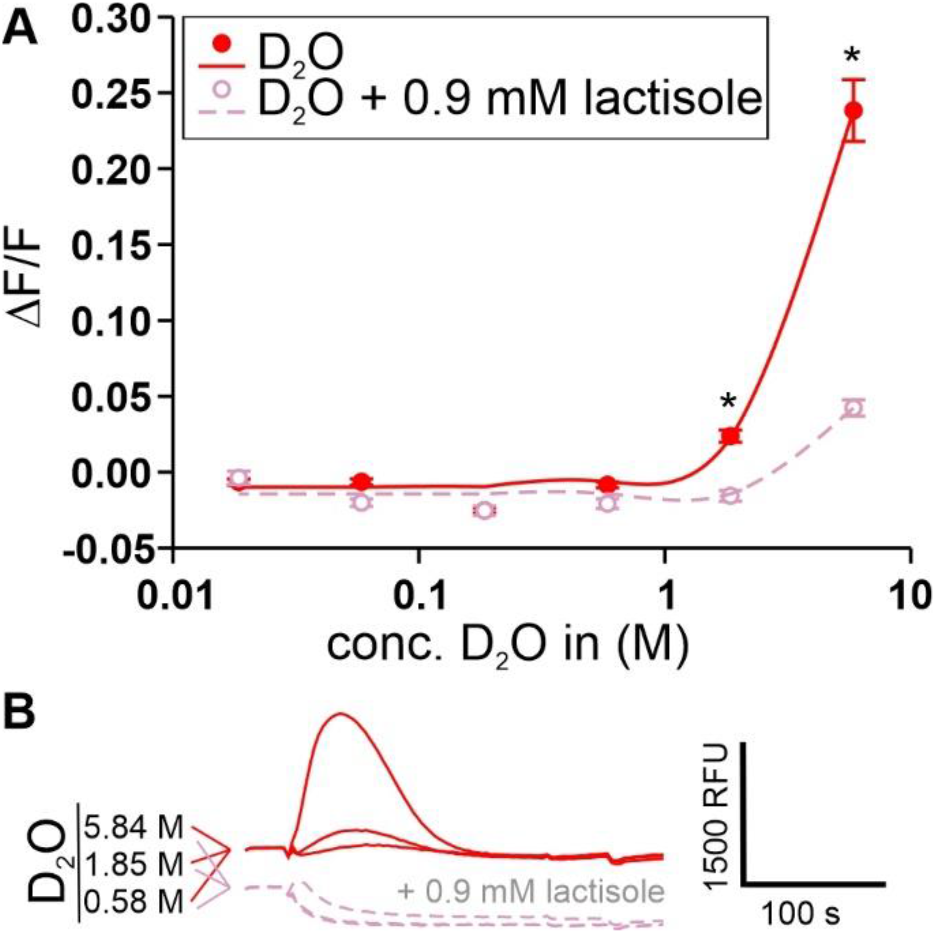
D_2_O-activation of the human sweet taste receptor. A) Dose-response relationship of cells expressing the human sweet taste receptor and treated with different concentrations of D_2_O (filled red circles, red line). Cells treated with lactisole served as negative controls (open pink circles, pink line). y-axis, relative changes in fluorescence upon stimulus application (ΔF/F). x-axis, logarithmically scaled molar D_2_O-concentrations. Asterisks indicate fluorescence changes above baseline significantly different from lactisole-treated controls (p ≤ 0.01). B) Raw fluorescence traces of D_2_O-treated (red-traces, top) and D_2_O + 0.9 mM lactisole-treated cells (pink-traces, bottom) stimulated with the indicated D_2_O-concentrations. A scale bar indicating relative fluorescence (relative fluorescence units (RFU) and experimental time (in seconds (s)) is included.

We further used an IP1 assay(*47, 48*) on non-transfected HEK293T cells, where we observed that dose-dependent curves of carbachol – an agonist of the endogenous muscarinic receptor 3 (M3)(*49*) – did not show any difference between H_2_O and D_2_O-based media (Figure 6A) and that cell medium that had either 10 % or 100 % D_2_O, did not activate basal IP1 accumulation (Figure 6B). Next, TAS1R2/TAS1R3 receptor along with the chimeric Gα16gust44 subunit(*46, 50*) were transiently expressed, and the functionality was illustrated by dose-dependent response to D-glucose (Figure 6C). Finally, and in agreement with calcium imaging, we found that 10 % D_2_O activated these cells. Activation by 100 % D_2_O was even more pronounced (Figure 6D).

**Figure 5.**
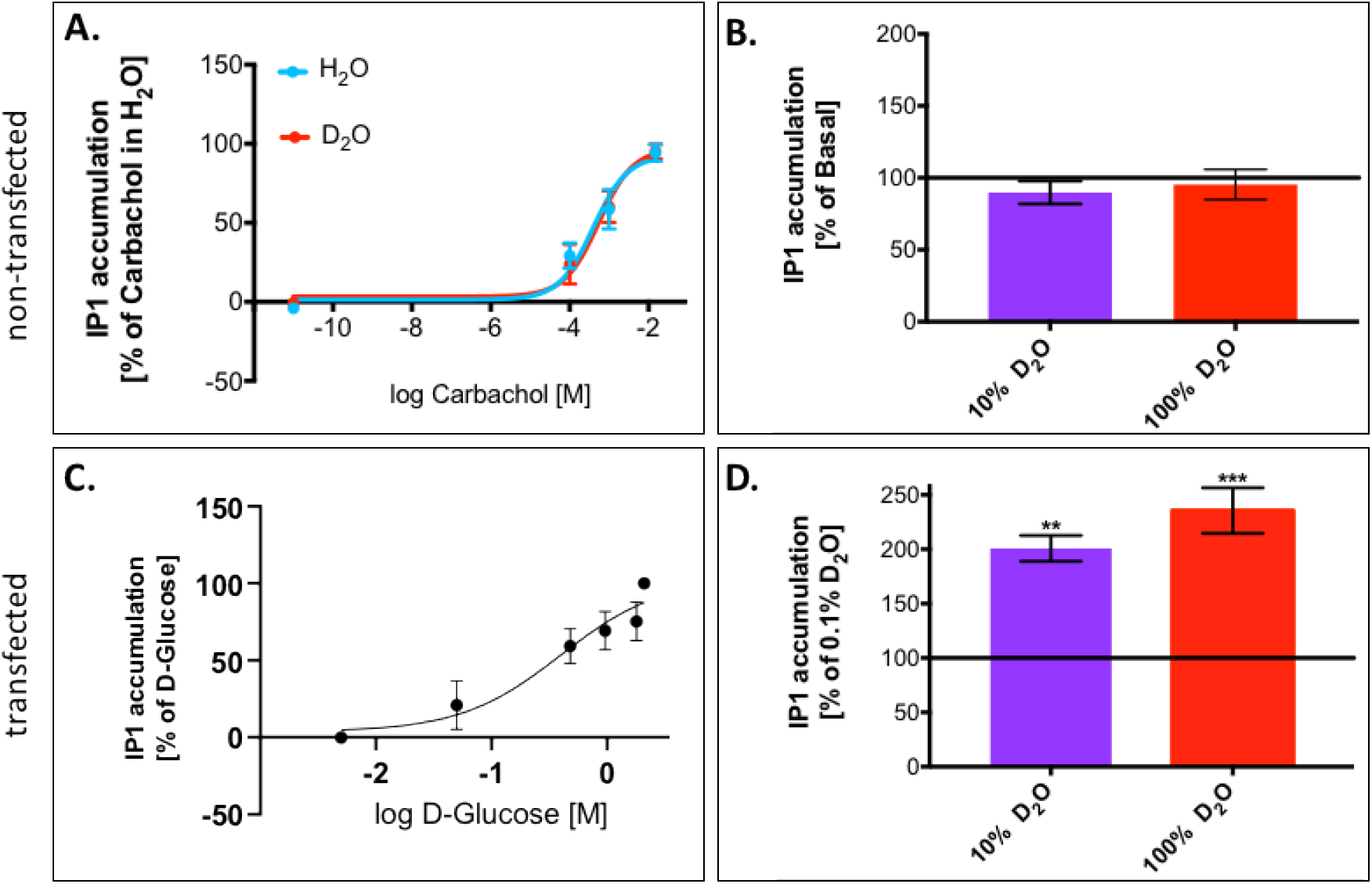
IP1 accumulation in HEK 293T cells following exposure to different ligands dissolved in powder-based DMEM medium. (A) Non-transfected HEK 293T cells respond similarly to raising concentrations of carbachol dissolved in D_2_O (red) as in H_2_O (blue). (B) D_2_O caused no elevation of IP1 levels in non-transfected HEK 293T cells. (C) HEK 293T cells transiently expressing TAS1R2/TAS1R3 respond positively to D-glucose. (D) Transfected HEK 293T cells are activated by D_2_O. Values represent the mean ± SEM of at least 3 replicates. The horizontal black line represents the basal values of controls. Significant differences in IP1 values from control values are marked with ** for p ≤ 0.005 and *** for p ≤0.0005 using Dunnett’s multiple comparisons test.

## Molecular modelling

The cellular response results further support the hypothesis that the sweet taste of D_2_O is mediated via the TAS1R2/TAS1R3 receptor. Various mechanisms governing this effect can be envisioned. As a potential suspect, we focus on a direct effect on the sweet taste receptor, narrowing on the TAS1R3 TMD (see Figure 1), as it is already known to be a modulation site with functional differences between humans and rodents(*23, 29, 30*). Furthermore, water-binding sites were discovered at the TMD of many GPCRs(*51, 52*), suggesting a potential target for D_2_O binding. We modeled the human TAS1R3 TMD using the I-TASSER server(*31*). Positions of H_2_O molecules were compared among mGluR5 structures (PDB: 4OO9, 5CGC, and 5CGD) and two conserved positions were found. The H_2_O molecules in these two positions were merged with the TAS1R3 model and minimized (Figure 7A). The water mapping protocol from OpenEye(*53*) enables mapping of water positions based on the energetics of water, and ~40 water molecules were predicted in the binding site using this protocol (Figure 7A). Water densities of H_2_O and D_2_O in the TMD of the TAS1R2/TAS1R3 receptor were calculated from MD simulations as described below. Overall, all three methods suggest the possibility for at least some internal molecules (trapped in the TMD bundle) in addition to water that surrounds the extracellular and intracellular loops (Figure 7A).

**Figure 7:**
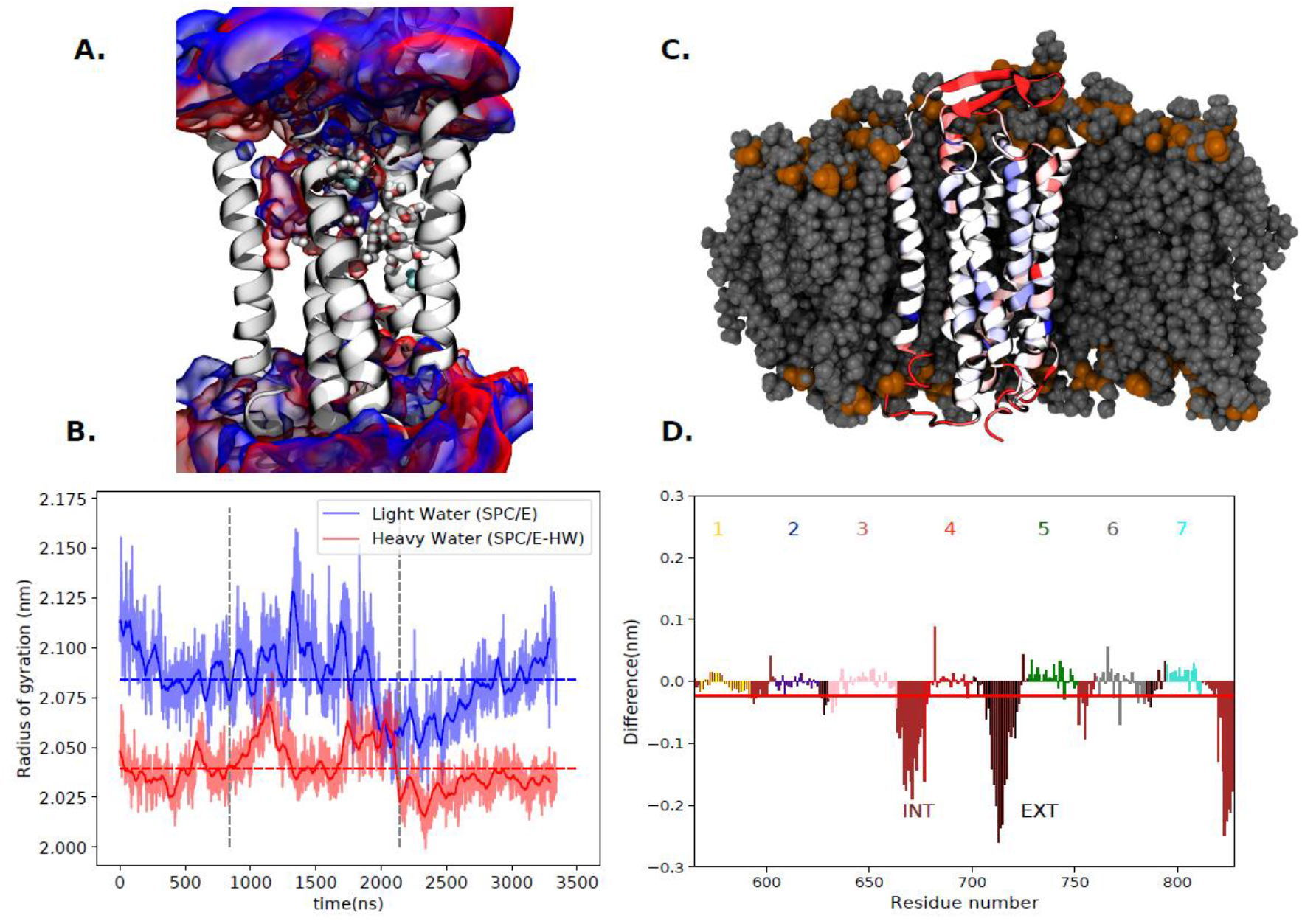
Differences between the behavior of the trans-membrane part of the human sweet taste receptor in H_2_O vs D_2_O base on analysis of three independent microsecond trajectories. (A) Structure of the TMD of the TAS1R2/TAS1R3 receptor with the probability density (volumetric map) of H_2_O (blue) or D_2_O (red) molecules within 10 Å from the protein evaluated using the VMD VolMap tool from the MD simulations at an isovalue of 0.1. The conserved water molecules in the x-ray templates are shown in cyan color. Water molecules predicted with the software OpenEye(*53*) are shown in licorice representation. (B) Time evolution of the radii of gyration in H_2_O (blue) and D_2_O (red) from three microsecond-timescale simulations (separated by vertical dashed lines) with total mean values as dashed lines, showing that the protein is more compact in heavy water. (C) Representative snapshot of the trans-membrane part of the human sweet taste receptor color-coded that red/blue represents parts more/less rigid in D_2_O vs H_2_O. The embedding lipid membrane is represented in gray. (D) Difference in root mean square fluctuations in MD trajectories. Negative/positive values mean that structures are more/less rigid in D_2_O than in H_2_O. The red line represents the sum over all residues.

Next, we carried out microsecond MD simulations of the TMD embedded in a phospatidylcholine (POPC) bilayer in either H_2_O or D_2_O (for details including our model of D_2_O effectively including nuclear quantum effects see SM). Note that water molecules enter the TMD domain and cluster at positions that partially overlap with the modeled water positions, see Figure 7A. More precisely, H_2_O and D_2_O have mutually slightly shifted densities inside the protein cavity, with H_2_O overlapping better than D_2_O with the modeled water positions. Furthermore, MD simulations show clustered water molecules close to the lactisole binding site. These internal positions may have a differential effect between H_2_O and D_2_O, though differences between the averaged water densities are not very pronounced. Figure 7B shows the time evolution of the radius of gyration of the TMD domain, while Figures 7C and 7D presents the root mean square fluctuations (RMSF) of individual residues of the proteins superimposed on its structure and plotted in a graph together with the mean value of RMSF. A small but significant difference is apparent in the behavior of the protein in H_2_O vs D_2_O. Namely, structural fluctuations of most residues (particularly those directly exposed to the aqueous environment) and of the protein as a whole are slightly attenuated in D_2_O, in which environment the protein is also somewhat more compact than in H_2_O (Figure 7B). Additional simulations on other representative systems show that the rigidifying effect of heavy water is apparent also in small soluble proteins (see SM).

## Summary and outlook

In summary, we have systematically addressed the question of the sweet taste of heavy water. Importantly, by employing gas chromatography/mass spectrometry analysis we demonstrate that the effect is not due to impurities. Being only isotopically different from H_2_O, in principle, D_2_O should be indistinguishable from H_2_O with regard to taste, namely it should have no taste of its own. H_2_O was shown previously to elicit sweet taste by rinsing sweet taste inhibitors away, both in human sensory experiments and in cell-based studies, which was explained in terms of a two-state model, where the receptor shifts to its activated state when released from inhibition by rinsing with water (*45*). Here, we have studied the taste of D_2_O and H_2_O *per se*, not related to washing away of sweet taste inhibitors. Using psychophysics protocols, we show that humans differentiate between D_2_O and H_2_O based on taste. Next, we illustrate that human subjects consistently perceive D_2_O as being mildly sweet and significantly sweeter than H_2_O. Moreover, D_2_O added to sweetness of some sweeteners; sweetness of glucose and cyclamate appears to be directly additive, while in the case of sucrose the additive effect was observed only at 50 mM sugar concentration. Furthermore, D_2_O did not enhance the umami taste perception of MSG, and reduced the perceived bitterness of 0.1mM quinine, in agreement with the known effect of bitterness suppression by sweet molecules.

A further important funding is that lactisole, which is an established blocker of the TAS1R2/TAS1R3 sweet taste receptor that acts at the TAS1R3 transmembrane domain(*30*), suppresses both the sweet perception of D_2_O in sensory tests and the activation of TAS1R/TAS1R3 in calcium imaging assay. In support of these observations cell-based experiments demonstrate that HEK 293T cells transfected with TAS1R2/TAS1R3 and Gα16gust44 chimera, but not the non-transfected cells, are activated by D_2_O, as measured by IP1 accumulation compared to control values. Finally, taste experiments on mice show that these animals do not prefer D_2_O over H_2_O.

Our findings point to the human sweet taste receptor TAS1R2/TAS1R3 as being essential for sweetness of D_2_O. Molecular dynamics simulations show, in agreement with experiment(*38*), that proteins in general are slightly more rigid and compact in D_2_O than in H_2_O. At a molecular level, this general behavior may be traced back to the slightly stronger hydrogen bonding in D_2_O vs H_2_O, which is due to a nuclear quantum effect, namely difference in zero-point energy (*3, 4*). Biologically relevant situations where one may expect strong nuclear quantum effects as implications of H/D substitution directly involve proton or deuteron transfer (*9*). Unless a yet unknown indirect mechanism is involved, this is not the case for the TAS1R2/TAS1R3 sweet taste receptor, thus the nuclear quantum effect is probably weak in the present case. Future studies should be able to elucidate the precise sites and mechanisms of action, as well as the reason why D_2_O activates TAS1R2/TAS1R3 in particular, resulting in sweet (but not other) taste. To this end, site directed mutagenesis as well as determination of the precise structure of the TAS1R2/TAS1R3 receptor will be of a key importance.

## Supporting information

SupportingInfo including methods and additional figures

## Acknowledgment

We thank R. F. Margolskee for the pcDNA of chimeric Gα16gust44 and P. Jiang and Y. Wang for hosting N.B.A. in Monell Chemical Senses Center during preliminary stages of this project. V.P. and L.M. acknowledge Ondřej Pačes and his team for the development of automatic drinking monitoring system, and Hedvika Vysušilová for outstanding technical assistance and animal handling. M.B. thanks Catherine Delaporte for excellent technical assistance. P.J. thanks the European Regional Development Fund OP RDE (project ChemBioDrug no. CZ.02.1.01/0.0/0.0/16_019/0000729) for support. Funding by ISF grant #1129/19 and UHJ-France and the Foundation Scopus to M.Y.N. is gratefully acknowledged. M.Y.N. is a member of COST actions Mu.Ta.Lig (CA15135) and ERNEST (CA18133). P.E.M. acknowledges support from his popular science YouTube channel.

## Author contributions

NBA, PEM, MYN, and PJ designed the study, NBA and HK performed sensory experiments, NBA and EM carried out IP1 experiments, MB conducted calcium imaging experiments, ND and YBSG performed homology modeling and structural analyses, PEM performed water purification, JC performed water purity tests, VP and LM performed experiments on mice, VCC developed the heavy water interaction potential, and CT performed the molecular dynamics simulations. PJ, MYN, and NBA wrote the paper with input from all coauthors.

## Notes

### Competing Interest Statement

The authors have declared no competing interest.

### Summary of Updates

Calcium imaging experiments have been added as Figure 5. Results, methods, abstract and outlook were edited to reflect this addition, which further supports the role of TAS1R2/TAS1R3 in mediating sweet taste of heavy water.

