## SupportingInfo including methods and additional figures for "Sweet taste of heavy water"

*[1]The Institute of Biochemistry, Food Science and Nutrition, The Robert H Smith Faculty of Agriculture, Food and Environment, 76100 Rehovot, Israel and The Fritz Haber Center for Molecular Dynamics, The Hebrew University of Jerusalem, 91904 Jerusalem, Israel, [2]Institute of Organic Chemistry and Biochemistry, Czech Academy of Sciences, Flemingovo nám. 2, 16610 Prague 6, Czech Republic, and [3]Leibniz-Institute for Food Systems Biology at the Technical University of Munich, Lise-Meitner-Str. 34, 85354 Freising, Germany.*

*[+]Both authors contributed equally.*

### Methods

#### *Sensory evaluation experiments*

A human sensory panel was used to resolve the gustatory effect in perception of $D_2O$ taste. Subjects between the ages 20 and 43 years were recruited. The study included 10 experiments with different groups of participants (15-30 subjects; between 4 to 12 males). The perception was tested by varying sensory tests as detailed below. Either sterile syringes with solutions (0.3 ml) or identical cups with solutions (7 ml), were presented in randomized order, unless otherwise noted. Participants were required to taste each solution using either 'tip of the tongue' or 'sip and spit' procedures, rinse their mouth with water after each solution and to wait for 30 seconds before moving to the next taste sample. All research procedures were ethically approved by the Committee for the Use of Human Subjects in Research in The Robert H. Smith Faculty of Agriculture Food and Environment, the Hebrew University of Jerusalem.

*Taste Solutions and concentration data:* 99.9 % purity $D_2O$ was purchased from Sigma-Aldrich Corp, while 18MΩ ultrapure grade was used for pure $H_2O$. All water samples were placed under vacuum and treated by ultrasound to remove any dissolved gases. See below for more details on water purification.

D-glucose (CAS Number: 50-99-7), sucrose (CAS Number: 57-50-1), cyclamate (CAS Number: 139-05-9), quinine (CAS Number: 207671-44-1) and MSG (CAS Number: 142-47-2) were purchased from Sigma-Aldrich Corp. All compounds were dissolved in both types of water to a final concentration of 50, 75 and 100 mM for D-glucose and sucrose; 3.5, 4.5 and 5.5 mM for cyclamate; 0.1 and 0.32 mM for quinine; 10, 25 and 50 mM for MSG. Lactisole (CAS Number: 150436-68-3) was purchased from Domino Specialty Ingredients and dissolved to a final concentration of 0.9 mM in $D_2O$ water as well in $H_2O$. The concentration of lactisole was selected based on previous data(*45, 54*). Sweeteners concentrations were selected to be in low intensity of sweetness. All solutions were prepared in the morning of the day of the experiment and were stored in individual plastic syringes (1 ml) for each participant.

*9-point scale (Figure 2):* The sweetness of heavy water and its effect on other taste compounds was evaluated in several independent experiments: (1) $D_2O$ sweetness relative to $H_2O$ (Figure 2A); (2) $D_2O$ effect on sweetness of D-glucose (Figure 2B), sucrose (Figure 2C) and cyclamate (Figure 2D); (3) $D_2O$ effect on quinine (Figure 2E) and MSG (Figure 2F); (4) Lactisole effect on $D_2O$ sweetness (Figure 4B); Intensity of each taste modality – sweetness/bitterness/umami was evaluated on a 9-point scale on Compusense Cloud, ranging from 1 (no sensation) to 9 (extremely strong sensation). In addition, participants had to report any additional tastes they recognized. Statistical tests were conducted using JMP Pro 13 (JMP, Version 13. SAS Institute Inc., Cary, NC, 1989-2019). Data were first analyzed employing ANOVA with participants as a random effect(*55*) . Thereafter the Tuckey Kramer test was used to compare mean sweetness between all samples(*55*). Significance was set at $p<0.05$, and preplanned comparison t-tests were used where relevant.

*Two-Alternative Forced Choice test*(*56*) *(2AFC, Figure 4A):* Participants were presented with two blind coded water samples of $D_2O$ and $D_2O$ + 0.9mM lactisole. The participants were asked to choose the sweeter solution. For the data analysis, the highest number of responses for one sample was compared to a statistical table(*56*) which states the minimum number of responses required for a significant difference.

*Testing of detectable difference between two samples with a triangle test*: Panelists were presented with two identical and one different water samples. All three samples were presented to the subjects at once, and the panelists were instructed to taste or smell the samples from left to right and identify the odd sample. Triangle tests were used to examine the difference in taste, as well as in smell, between $H_2O$ and $D_2O$. Participants tasted each solution from individual plastic syringes and the procedure was repeated twice, without ('taste and smell' Figure S1A) and with ('taste only', Figure S1B) nose clips. The 'taste only' experiment included 26 participants (5 males) and the 'smell and taste' experiment included 28 subjects (9 males). In examining the effect of smell (Figure S1C), the experiment included 25 subjects (6 males) who were asked to smell each sample in a glass jar(5ml). For the data analysis, the total number of responses correctly identifying the 'odd' sample was counted and compared to a statistical table which determines the critical number (minimum) of correct answers required for a significant difference.

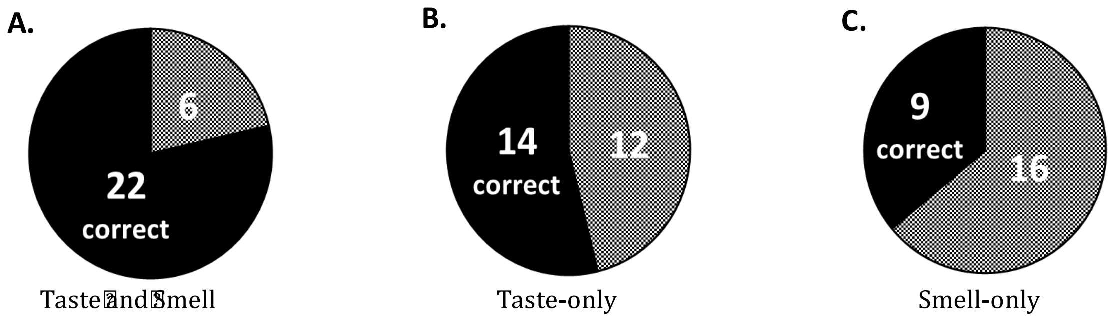


**Figure S1.** Discrimination tests of $D_2O$. $D_2O$ can be distinguished from $H_2O$ based on taste. Numbers within a circle indicate the number of subjects that correctly (black) or incorrectly (grey) identified the odd solution.

***Details of the water purification procedure***

Water purification method was designed to achieve two things: i) to remove all volatile components from water and ii) remove all non-volatile components. An all borosilicate glass apparatus with a 250 ml flask and a side arm with five takeoff points for ampoules was created. The apparatus was completely washed three times with 18 MΩ Millipore water and dried under vacuum. The glassware was flamed under vacuum to 400 C. This has the effect of burning off any volatiles in the apparatus. The kit was then charged with 180 ml of $H_2O$ (18 MΩ Millipore) or $D_2O$ (Sigma-Aldrich, 99.9 %). Water was then pumped out under vacuum to a teflon headed pump for 10 minutes while simultaneously being sonicated, which removes any dissolved gas. The flask was then heated to 60 °C. Water was then 'distilled' (vapor transported) to the fifth ampoule submerged in iced water. After 5 ml of water were condensed, this ampoule was sealed with a glass torch and discarded. The remaining water was distilled into four 40 ml ampoules (using again iced water), each being sealed under vacuum with a glass torch. The remaining water in the apparatus was discarded. The same apparatus and procedure were used for both $H_2O$ and $D_2O$.

The purity of $H_2O$ and $D_2O$ was further checked by gas chromatography coupled to mass spectrometry (GC/MS). The organic impurities that may have been present in the water were extracted with hexane (distilled in a glass apparatus from analytical grade solvent supplied by Penta, Czech Republic). A sample of water (6 ml) was shaken vigorously for 5 minutes with 2 ml of hexane in a ground-glass stoppered test tube. The organic layer was collected and its volume was reduced to 200 μl under a stream of nitrogen. The analyses were performed on a 6890N gas chromatograph coupled to a 5975B quadrupole mass spectrometer (Agilent Technologies, Santa Clara, CA). The sample (1 μl) was injected in the split mode with a split ratio of 10:1 and with the injector temperature set to 200 °C. An HP-5MS fused silica capillary column (30 m × 250 μm; a film thickness of 0.25 μm) from Agilent Technologies was used for the chromatographic separation. The carrier gas was helium at a constant flow rate of 1.0 ml/min and the temperature program was set as follows: 40 °C (2 min), then 8°C/min to 200°C (0 min), then 15°C/min to 320°C (3 min). The transfer line, ion source, and quadrupole temperatures were set to 280 °C, 230°C, and 150°C, respectively. The ionization was achieved using 70 eV electrons and the solvent delay time of 4 min was used. The total ion current chromatograms (*m/z* 29 - 600 range) shown in Figure S2 displayed background, without any signs of chromatographic peaks. Therefore, it can be concluded that the water samples used for the experiments did not contain significant amounts of volatile or semi-volatile organic compounds.

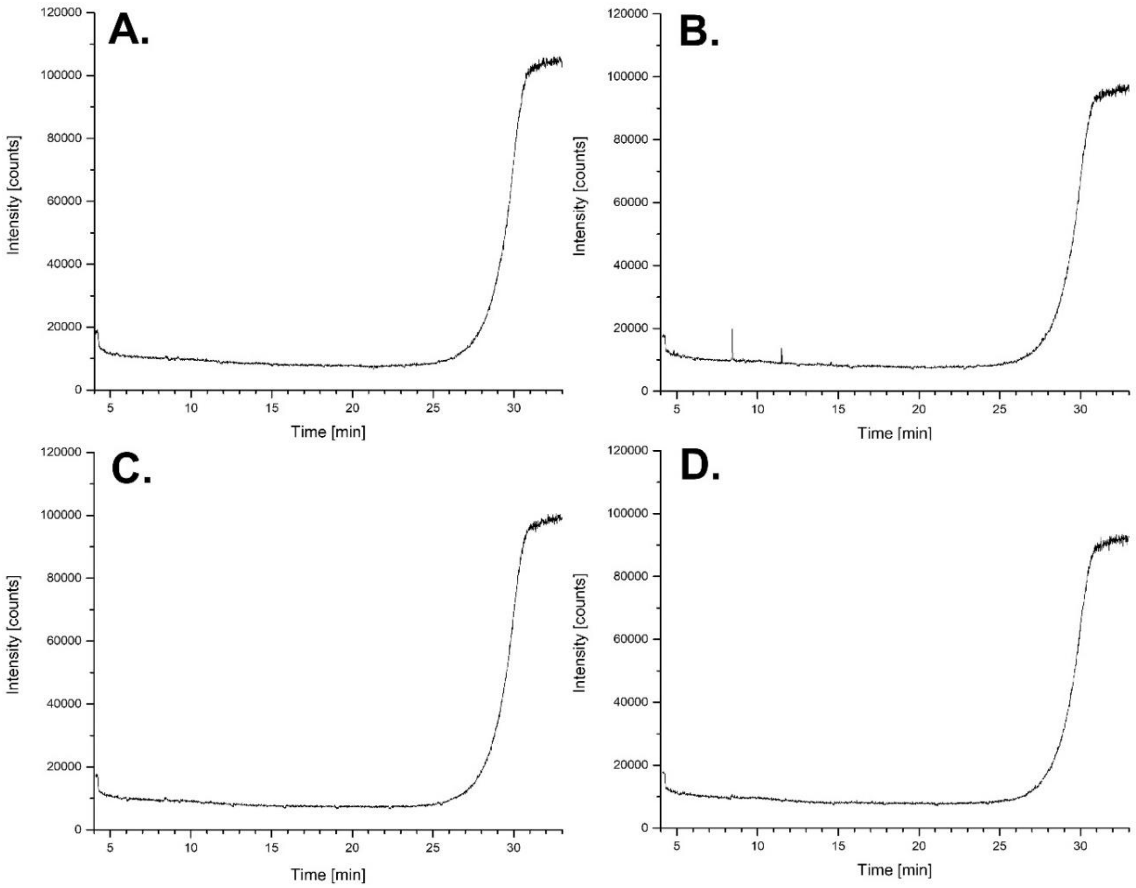


**Figure S2**. GC/MS chromatograms of hexane extracts of $H_2O$ (A., B.) and $D_2O$ (C., D.) showing clean background, without any chromatographic peaks. The background signals have typical profiles, with a tail of the solvent peak (4 min) and a rising signal at high retention times due to stationary phase bleeding at elevated temperatures. Small signals in panel B. are not chromatographic peaks but spikes caused by air occasionally penetrating the detector through the instrument fittings.

#### *Ruling out impurity effects on taste*

As seen in Figure S3, $D_2O$ was rated sweeter than $H_2O$ for all $D_2O$ and $H_2O$ purities that were tested, i.e., from the first, second, third, and fourth distillation batch. Here we used the labeled magnitude scale (LMS) for sweetness evaluation. Each participant (n=28; 8 males) was asked to taste eight solutions, four per each type of water, which differed in their level of purity. The solutions were offered in order of ascending purity and randomized in terms of $D_2O$ and $H_2O$. The scale was bound by 'no sensation' at the bottom and 'strongest imaginable' at the top of the range(*57*).

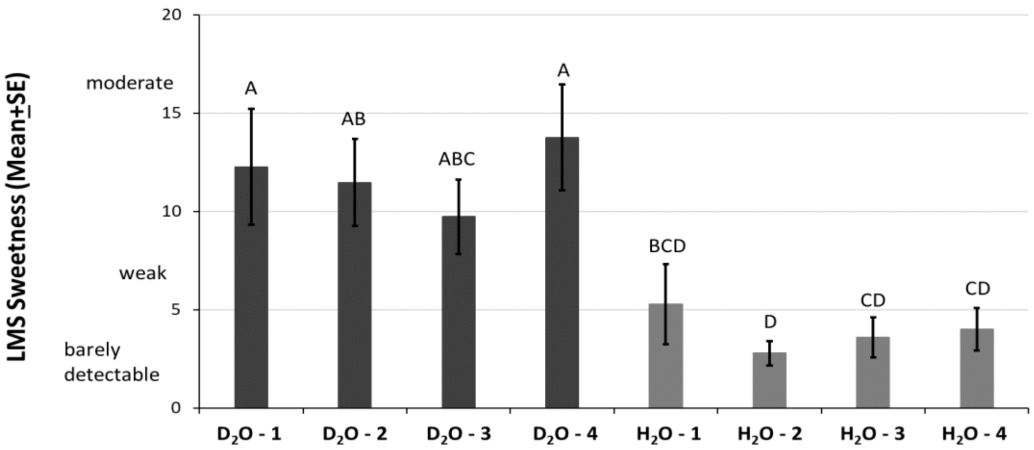


**Figure S3.** Difference in sweetness between two types of water at the same level of purity. Statistical analysis was performed using the two-way analysis of variance (ANOVA) with a Tuckey Kramer test (n=30; 9 males); samples not connected by the same letter are significantly different ($p<0.05$). The data are presented as the mean ± the standard error of measurement (SEM). The y axis shows the response for sweetness on LMS scale, and the x axis is labeled with different water samples.

#### *Heterologous Expression*

*IP-one assay* - TAS1R2/TAS1R3 stimulated activation of the G protein mediated pathway was measurement applying the IP-One HTRF assay (Cisbio) based on the manufacturer's protocol. In brief, HEK 293T cells (ATCC) were grown to a confluency of approximately 85-90 % and transiently transfected with 6μg/plate DNA (TAS1R2, TAS1R3, Gα16gust44) by applying LipofectamineTM 2000 (Invitrogen, USA, 30μl/plate) transfection reagent, according to the manufacturer's protocol. The next day, cells were suspended with fresh Dulbecco's Modified Eagle's (DMEM) Medium, containing 10% Fetal Bovine Serum (FBS), 1% L- glutamine amino acid and 1% penicillin streptomycin (10% DMEM), seeded (0.5 ml cells per well) into 24-well culture plate, and maintained for 8-12 h at 37 °C. Then cells were "starved" overnight by changing the medium to 0.1% DMEM (containing 0.1% FBS), in order to reduce the basal activity of the cells. Cells exposure was performed by addition of 0.5 ml tested compound dissolved in 0.1 % DMEM with 50 mM Lithium Chloride (LiCl) for 5 minutes directly into the wells. The presence of LiCl in this step is crucial because LiCl leads to IP1 accumulation(*47*). At the end of exposure time, tastant solution was replaced with fresh medium (0.1 % DMEM) containing 50 mM LiCl for another 55 minutes. Later, wells were washed with 100µl cold phosphate buffered saline (PBS) + Triton X-100, and kept at -80◦C for a few hours, in order to dissolve the cell membrane. For the IPOne HTRF assay, cell lysate was mixed with the detection reagents (IP1-d2 conjugate and Anti-IP1 cryptate TB conjugate, each dissolved in lysis buffer), and added to each well in a 384-well plate for 60min incubation at room temperature. Finally, the plate was read using Clariostar plate reader (BMG, Germany) equipped with 620 ± 10 nm and 670 ± 10 nm filters. IP1 levels were measured by calculating the 665nm/620nm emission ratio. All responses are presented as the means ± SEM of IP1 accumulation (%). Dose–response curves were fitted by non-linear regression using the algorithms of PRISM 7 (GraphPad Software, San Diego, CA, USA). Column figures were analyzed using one-way ANOVA with a Dunnett's (*55*). Each compound was tested in triplicate in three individual experiment in comparison to the reference (carbachol dissolved in $H_2O$ or basal levels)(*47*).

*Compounds* - In the case of $D_2O$, in order to test its specific effect, we used a powder DMEM medium (CAS Number: D5030, Sigma Aldrich), dissolved in the needed amount of $D_2O$ instead of the liquid one. Other ligands than $D_2O$ were purchased from Sigma-Aldrich or Domino Specialty Ingredients as noted in ‘Taste Solutions and concentration data' above. Unless noted otherwise, ligands were used at final concentrations of 5, 50, 480, 960, $1.8 \times 10^3$ and $2.1 \times 10^3$ mM for D-glucose; 0.1, 1 and 15 mM for carbachol and $D_2O$ at 0-100 % proportionately to $H_2O$. Solutions were at pH=7.4.

*Calcium mobilization assay* - For the functional assays with the human sweet taste receptor we used a cell line (HEK 293 FlpIn T-Rex), which constitutively expresses the sweet taste receptor subunit TAS1R2 as well as the chimeric G protein Gα15gi3, whereas the sweet taste receptor subunit TAS1R3 can be induced by tetracycline(*45, 46*). The functional experiments were done as described before(*46*). Briefly, the cells were grown in DMEM supplemented with 10% fetal bovine serum, 100 U Penicillin/mL, 0.1 mg/mL Streptomycin, 2 mM L-glutamine, at 37°C and 5%-$CO_2$, 100% air humidity. The day before the experiment, cells were seeded to a density of 50-60% onto 96-well plates coated with 10 µg/mL poly-D-lysine and 0.5 µg/mL tetracycline was added. Next, cells were loaded with Fluo-4 AM in the presence of 2.5 mM probenecid for 1 h. After this, cells were washed twice with C1-buffer (130 mM NaCl, 5 mM KCl, 10 mM HEPES, 1 mM sodium pyruvate, and 2 mM $CaCl_2$, pH 7.4) before placing them in a fluorometric imaging plate reader (FLIPR$^{tetra}$, Molecular Devices) for measurements.

C1-buffer prepared with $D_2O$ was mixed with C1-buffer made with $H_2O$ to result in the following final $D_2O$-concentrations (a further 3-fold dilution, which occurs upon application of 50 µL stimulus to 100 µL of C1-buffer in the 96-well plates is already included): 18.47 M, 5.84 M, 1.85 M, 0.584 M, 0.185 M, 0.058 M, 0.018 M, 0.000 M. Fluorescence changes were monitored after automated application of stimuli. As specificity control C1-$D_2O$ including 0.9 mM lactisole, a selective inhibitor of the human sweet taste receptor(*58*), was applied to identically treated cells. Experimental results from five biological replicates performed in quadruplicates were used to establish the dose-response relationship using the software SigmaPlot as before(*46*). As the highest $D_2O$-concentration resulted in fluorescence changes largely resistant to lactisole blocking, the 18.47 M concentration was excluded. Student's t-test was used to confirm that $D_2O$-induced fluorescence changes above baseline were significantly ($p \leq 0.01$) different from lactisole-treated controls.

***Animal experiments***

All animal experiments followed the ethical guidelines for animal experiments and the Act of the Czech Republic Nr. 246/1992 and were approved by the Committee for Experiments with Laboratory Animals of the Czech Academy of Sciences. Three-month-old male C57BL/6J mice (n = 34) from Charles Rivers Laboratories (Sulzfeld, Germany) were housed at a temperature of 23 °C with a daily cycle of 12 h light and dark (lights on at 6 am). The mice were placed in groups of two in cages with automatic drinking monitoring system (Developmental Workshops of Institute of Organic Chemistry and Biochemistry of the Czech Academy of Sciences, Prague, Czech Republic). They were given *ad libitum* water and a standard rodent chow diet (Ssniff Spezialdiäten GmbH, Soest, Germany). On the day of the experiment, during the dark phase of the cycle, freely fed mice were given weighed food pellets and two 30 ml glass bottles. The two bottles contained pure $H_2O$ and pure $D_2O$ (n = 12), or $H_2O$ and sucrose solution (43 mmol/l sucrose solution in $H_2O$) (n = 10), see Table S1. Mice drinking $H_2O$ in both bottles served as a control group (n = 12). Drinking was monitored every 10 min for 16 h (starting from 6 pm) and food intake was determined at the end of the experiment (Figure S4).

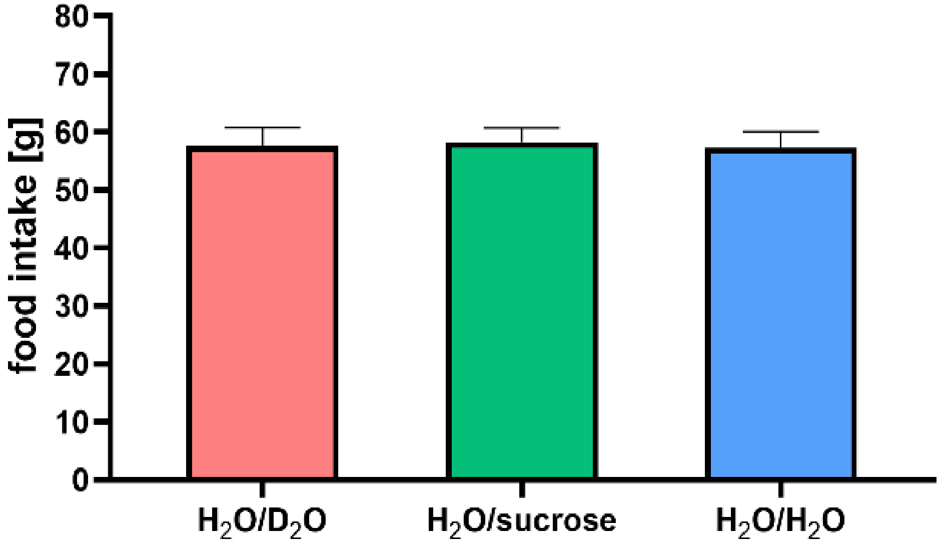


**Figure S4.** Food intake during experiment. Mice were placed in groups of two in cages. The data are presented as the mean ± standard error of measurement (SEM). Statistical analysis was performed using the one-way ANOVA with a Dunnett's test. (n = 10-12)

All responses are presented as the means ± SEM. Statistical analysis was performed using ANOVA with a Dunnett's test(*55*) for food intake. Two-way ANOVA with a Bonferroni's multiple comparisons test was used for analysis of average volume of liquid consumption. Analysis was performed using the GraphPad Software, Inc., Prism 8 (GraphPad Software, San Diego, CA, USA). The differences between the control and treated groups were considered significant at $p < 0.05$.

**Table S1.** Design of the animal experiment.

| condition 1 | condition 2 | control |
|---|---|---|
| $H_2O$ | $H_2O$ | $H_2O$ |
| $D_2O$ | sucrose | $H_2O$ |

### Modeling and Docking

All models were prepared with I-Tasser web server(*31*). The templates that were used by I-Tasser for each of the TAS1R2 and TAS1R3 monomers were: 6N51, 5X2M, and 5K5S. To model the full heterodimer, the monomer models were aligned to a Class-C GPCR Cryo-EM structure (PDB: 6N51) and minimized with Schrödinger Maestro 2019-1. To illustrate the orthosteric binding site of sugars D-Glucose was prepared (Schrödinger Maestro 2019-1, LigPrep) and docked with Glide XP , to the TAS1R2 VFT domain, model that was based on 5X2M in a protocol that was validated in previous work(*32*). The TAS1R3 binding site is based on a lactisole molecule docked to TAS1R3 TMD model (Schrödinger Maestro 2018-2, Glide SP, template PDB ID: 4OR2 and 4OO9). The figure was made using ChimeraX (version 0.93)(*33*). Water molecules were predicted with a water mapping software (SZMAP 1.5.0.2: OpenEye(*53*)) after a successful benchmark over conserved water templates (PDB IDs 5CGC and 5CGD), in which the software was able to identify overall 8 out of 10 crystal water molecules around the ligands of the templates.

#### *Homology modeling and sequence comparison*

Several models of the transmembrane region of TAS1R3 and Tas1r3 were predicted by I-TASSER server. The best predicted model was chosen for further analysis, according to C-scores. mGluR1 and mGluR5 structures (PDB IDs: 4OR2 and 4OO9, respectively) were used as templates for constructing the models. Selected human and mouse models were then minimized and refined, using scwrl4(*59*) and Schrodinger program suits (version 11.2, Schrodinger, LLC, New York, NY, 2014).

Positions of $H_2O$ molecules were compared among mGluR5 structures (4OO9, 5CGC, and 5CGD) and two conserved positions were found in the area of the TMD of mGluR5 structures.

Docking was performed for cyclamate and lactisole, to both human and mouse TAS1R3, in order to elucidate putative interaction of these compounds on the allosteric binding active site of the TMD of TAS1R3 and Tas1r3. Both ligands could be docked at the allosteric pocket of TAS1R3, but neither cyclamate nor lactisole docked into Tas1r3 model. This is in accord with experimental data showing that mice are not sensitive to cyclamate and lactisole.

***Development of effective $D_2O$ model for simulations***

We have developed a new heavy water model SPC/E-HW (for intermolecular force field parameters of all the employed water models see Table S2) which effectively takes into account nuclear quantum effects (zero point motions in particular) by modifying the interaction potential for classical molecular dynamics. We were forced to develop a new model since the existing SPC/HW model (based on the same principles)(*60*) is not reproducing experimental properties of $D_2O$ very well (see comparison with the present model in Tables S3 and S4). The results presented in Tables S3 and S4 were obtained using MD simulations with a time step of 2 fs, a long-range interaction cut-off of 1.2 nm, long range corrections for energy and pressure, the Nose-Hoover thermostat(*61*) with a coupling constant of 1.0 ps, and the Parrinello-Rahman barostat(*62*) with a coupling constant of 2.5 ps and a compressibility of $5 \cdot 10^{-5}$ bar$^{-1}$. The viscosity of the water model was calculated by the Green-Kubo formula(*63*). All simulations were performed using the Gromacs program package(*64*), version 5.1.2.

**Table S2**: Oxygen Lennard-Jones parameters σ and ε and charges $q_O$ of the water models used in this paper. All of them use the SPC/E geometry where the distance between oxygen and hydrogen atoms ($d_{OH}$) is 1 and the angle between those atoms of 109.47°.

| Water model | σ/nm | ε/(kJ/mol) | $q_O/e^-$ |
|---|---|---|---|
| SPC/E-HW | 0.31970 | 0.5050 | 0.8376 |
| SPC/HW(*60*) | 0.31657 | 0.6497 | 0.8700 |
| SPC/E(*65*) | 0.31656 | 0.6502 | 0.8476 |

**Table S3**: Densities, temperatures of maximum density (TMD), and viscosities of the old (SPC/HW) and present (SPC/E-HW) heavy water models and the standard SPC/E normal water model. The experimental values are given in brackets.

| | Density $\rho/(kg^3/m)$ | TMD/K | Viscosity η/(mPa·s) |
|---|---|---|---|
| SPC/E-HW | 1104.0±0.3<br>(1104.4) | 250±1 (284.3) | 0.88<br>(1.097) |
| SPC/HW(*60*) | 1125.6±0.3<br>(1104.4) | 268±1 (284.3) | 1.40<br>(1.097) |
| SPC/E(*65*) | 999.2±0.4<br>(997.1) | 248±1 (277.1) | 0.73<br>(0.890) |

**Table S4**: Density differences, temperature of maximum density (TMD) differences, and viscosity differences between heavy and light water for the old (SPC/HW) and present (SPC/E-HW) heavy water models (with the standard SPC/E normal water model as a reference). Experimental values are in brackets.

| | $\Delta\rho/(kg/^3m)$ | $\Delta$TMD/K | $\delta\eta$(%) |
|---|---|---|---|
| SPC/HW(*60*) | 126.4<br>(107.3) | 20<br>(7.2) | 91.8<br>(23.3) |
| SPC/E-HW | 104.8<br>(107.3) | 2<br>(7.2) | 20.6<br>(23.3) |

***Molecular dynamics simulations***

The initial structure of the trans-membrane part of the human sweet taste receptor was designed by the homology modeling described above. It was then embedded in a membrane bilayer formed of 128 POPC lipids solvated either in $H_2O$ or $D_2O$, with chloride added to neutralize the systems, employing the CHARMM36 forcefield(*66*). For each water model a total of 5 µs has been run at 298K and 1 atm, with about 3 µs used for analysis. All simulations were performed using the Gromacs program package(*64*), version 5.1.2.

We checked robustness of the MD simulations for the TMD of TAS1R3 by generating three independent trajectories and plotting the observables for each one separately. As seen from Figures S5 (and Figure 7B in the main paper), our sampling is sufficient and both differences in root mean square fluctuations and the radii of gyration are reasonably converged. Figure S6 displays for three independent simulation runs the time evolutions of the root mean square deviations, from which the RMSF and differences thereof in $H_2O$ vs $D_2O$ were evaluated.

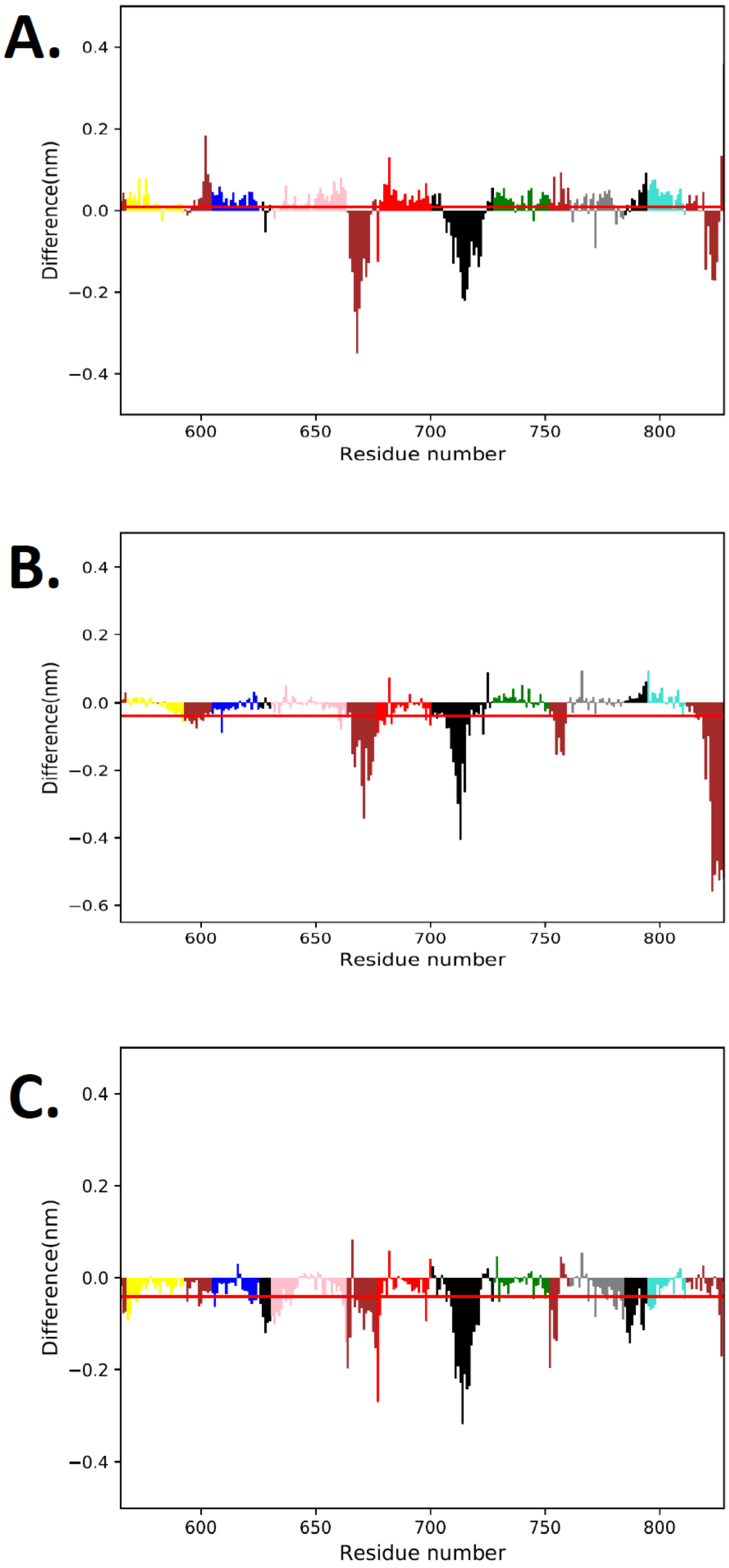


**Figure S5:** Differences in root mean square fluctuations of individual residues and their sum (red) for three separate microsecond-timescale simulations (color-coding is the same as in Figure 7D in the main paper).

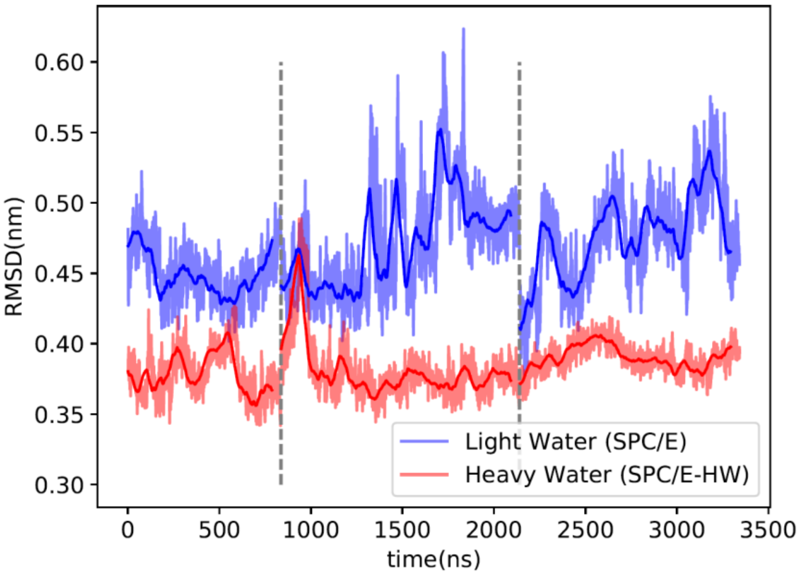


**Figure S6:** Time evolution of the RMSD in $H_2O$ vs $D_2O$ for three independent microsecond-timescale simulations (separated by vertical dashed lines).

In addition, we performed MD simulations of two other proteins - apo-azurin (PDB: 1E65) and ribonuclease-T1 (PDB:2RNT) – for which experimental data in $H_2O$ and $D_2O$ were available and the effect of heavy water on structural flexibility was studied(*38*). We employed the CHARMM36 forcefield(*66*). The proteins were solvated and $Na^+$ ions were added to zero the total charge. This system was energy-minimized and then simulations (two replicas) were run for 700 ns at 298K and 1 atm using the Nose-Hoover thermostat(*61*) and the Parrinello-Rahman barostat(*62*). For the time averaged analysis, the first 200 ns were discarded. We checked that the root mean square displacements from the initial structure did not drift in time and were relatively small (< 2 Å). All simulations were performed using the Gromacs program package(*64*), version 5.1.2.

From the MD simulations of the proteins in $H_2O$ (SPC/E[52]) and $D_2O$ (SPC/E-HW, see below) we calculated the time evolution of the radii of gyration. We see that most residues are slightly more rigid and the protein structures are a slightly more compact in $D_2O$ compared to $H_2O$ (Figure S7), which is consistent with the experimentally observed higher protein rigidity in the former solution(*38*).

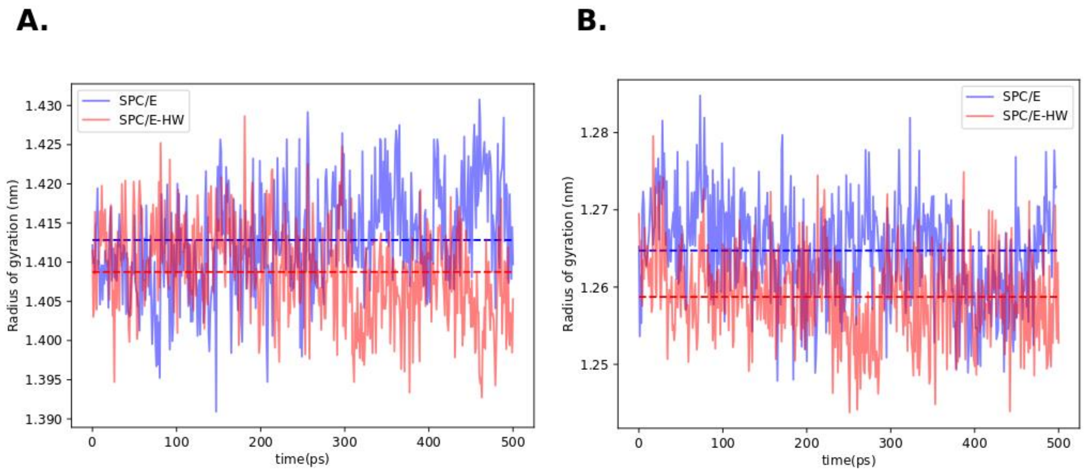


**Figure S7:** Time evolutions of the radii of gyration in $H_2O$ (blue) and $D_2O$ (red) with mean values as dashed lines, showing that both (a) apo-azurin and (b) ribonuclease-T1 are more compact in heavy water.
